# Frequency-Domain Analysis Links Autonomic Disruption to Renal Autoregulatory Failure after Spinal Cord Injury

**DOI:** 10.64898/2026.06.29.735393

**Authors:** Angela Tsang, Gagandeep Kaur, Veronica J. Tom, Evren Gurkan-Cavusoglu, Patrick Osei-Owusu

**Affiliations:** Department of Electrical, Computer and Systems Engineering, Case Western Reserve University School of Engineering, Cleveland, OH 44106; Department of Physiology & Biophysics, Case Western Reserve University School of Medicine, Cleveland, OH 44106; Department of Neurobiology & Anatomy, Drexel University College of Medicine, Philadelphia, PA 19129

**Keywords:** Spinal cord injury, renal hemodynamics, renal autoregulation, sympathetic nerve activity, myogenic mechanism, blood pressure variability

## Abstract

Spinal cord injury (SCI) disrupts supraspinal autonomic pathways that regulate cardiovascular function, producing marked blood pressure instability and contributing to secondary injury in peripheral organs. The kidney is particularly vulnerable to these disturbances because renal blood flow (RBF) depends on tightly regulated interactions between neural, myogenic, and vascular control mechanisms. However, how SCI level and chronicity alter dynamic renal autoregulation remains poorly defined. Here, we investigated the effects of high- and low-thoracic SCI on renal hemodynamic control using in vivo blood pressure and RBF recordings in female mice. Hemodynamics were assessed at baseline and during acute sympathetic stimulation induced by norepinephrine (NE; 10 µg/kg, i.v.) at 24 h and 4 wk following spinal cord transection at thoracic level 3 (T3) or thoracic level 10 (T10). Time-domain analyses quantified systolic blood pressure recovery, while frequency-domain analyses were used to resolve myogenic and sympathetic contributions to RBF regulation. High-thoracic SCI caused marked disruption of renal vascular responses to acute hypertension, producing paradoxical increases in RBF during NE-induced pressure elevations and sustained reductions in baseline and evoked RBF activity within frequency ranges associated with myogenic and sympathetic vasomotion. These impairments were most pronounced during the chronic phase of injury, consistent with loss of dynamic autoregulatory control and vascular remodeling. In contrast, low-thoracic SCI preserved baseline renal vasomotor activity and demonstrated recovery of dynamic autoregulatory responses over time. These findings identify SCI level and chronicity as critical determinants of renal microvascular regulation and demonstrate that high-thoracic SCI produces persistent autonomic-vascular uncoupling. This disruption of dynamic renal autoregulation represents a previously underappreciated mechanism of secondary organ vulnerability following neurotrauma.

**VISUAL ABSTRACT:** 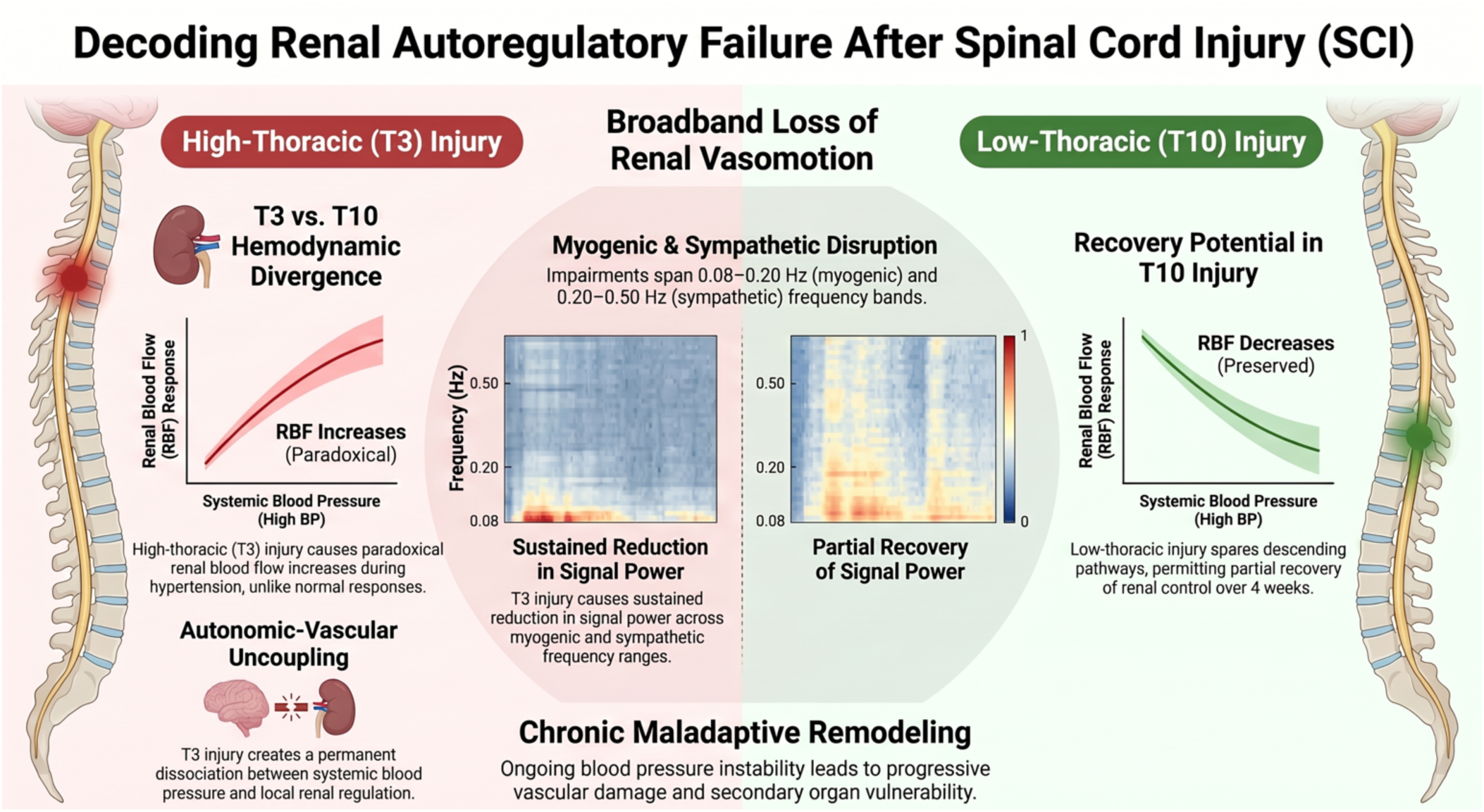

*New & Noteworthy:* This study introduces a frequency-domain analytical framework to quantify dynamic renal blood flow regulation after spinal cord injury (SCI). By resolving myogenic and sympathetic vasomotor activity across physiologically relevant frequency bands, we reveal that high-thoracic SCI causes a sustained, broadband loss of renal vasomotion and impaired autoregulatory responses to acute hypertension. In contrast, low-thoracic SCI preserves baseline spectral activity and permits recovery over time. This approach provides a mechanistic link between autonomic disruption and renal vascular dysfunction following SCI.

## INTRODUCTION

Spinal cord injury (SCI) is a devastating neurological disorder that results not only in loss of motor and sensory function but also in profound disruption of autonomic regulation. Injury-induced interruption of descending supraspinal pathways alters sympathetic and parasympathetic outflow, producing severe cardiovascular instability that contributes substantially to morbidity and mortality in individuals with SCI^1–3^. These disturbances manifest clinically as neurogenic shock, chronic hypotension, exaggerated blood pressure variability, and life-threatening episodes of autonomic dysreflexia, with severity strongly dependent on the anatomical level and completeness of the injury^4–6^.

Disruption of autonomic cardiovascular control following SCI has important implications for secondary injury in peripheral organs. Highly perfused visceral organs, including the kidney, are particularly vulnerable to the large and repeated fluctuations in systemic arterial pressure that accompany autonomic dysregulation after SCI^7,8^. Clinical and experimental studies have linked SCI to progressive renal dysfunction, acute kidney injury, and increased long-term risk of chronic kidney disease, suggesting that impaired neural control of renal perfusion represents a critical but underappreciated consequence of neurotrauma^7–9^.

Under physiological conditions, the kidney maintains relatively stable renal blood flow (RBF) and glomerular filtration rate across a wide range of perfusion pressures through highly efficient autoregulatory mechanisms^10–12^. This dynamic regulation is mediated by a rapid myogenic response of the afferent arteriole and a slower tubuloglomerular feedback mechanism, both of which act to buffer pressure fluctuations and protect the renal microvasculature^12–17^. In rodents, myogenic regulation predominates in low-frequency pressure oscillations (approximately 0.08–0.30 Hz), whereas sympathetic vasomotor activity contributes at higher frequencies (approximately 0.20–0.50 Hz), allowing frequency-domain analysis of RBF to resolve neural and vascular control components^17–20^.

Renal vascular tone is strongly influenced by central autonomic pathways originating in supraspinal nuclei that project to sympathetic preganglionic neurons within the thoracolumbar spinal cord^1,21^. High-thoracic SCI disrupts these pathways, functionally isolating the splanchnic and renal vascular beds from supraspinal sympathetic control, whereas lower-thoracic injuries spare critical descending projections and preserve a degree of autonomic regulation^1,6,21^. In the acute phase of high-level SCI, abrupt loss of sympathetic vasomotor tone produces neurogenic shock characterized by severe hypotension, bradycardia, and marked reductions in renal plasma flow and glomerular filtration^2,21^. Chronically, maladaptive plasticity within decentralized spinal autonomic circuits promotes exaggerated sympathetic reflexes and extreme blood pressure variability, particularly during episodes of autonomic dysreflexia^3,5,6,22^.

We previously demonstrated that chronic blood pressure instability associated with high-thoracic SCI produces severe structural and functional damage to the renal microvasculature, including loss of autoregulatory capacity and pathological extracellular matrix remodeling^23^. Rather than constricting in response to elevated perfusion pressure, the injured renal vasculature paradoxically dilates, exacerbating pressure transmission to downstream microvessels and promoting renal injury^23^. These findings suggest that SCI-induced autonomic disruption impairs dynamic renal autoregulatory mechanisms, particularly the myogenic response, but the temporal evolution and frequency-specific characteristics of this dysfunction remain incompletely understood.

In the present study, we tested the hypothesis that SCI level and chronicity differentially disrupt dynamic renal autoregulation through loss of supraspinal autonomic control. Using in vivo hemodynamic recordings in a mouse model of thoracic SCI, we combined time-domain analysis with frequency-domain signal processing to quantify renal vascular responses to acute sympathetic stimulation with norepinephrine. This approach enabled resolution of frequency-specific alterations in myogenic and sympathetic vasomotor activity and allowed us to define how neural injury level shapes renal microvascular control across acute and chronic phases of SCI. By linking autonomic disruption to impaired renal autoregulation, these findings provide new mechanistic insight into secondary organ vulnerability following neurotrauma.

## EXPERIMENTAL METHODS

The following is a summary of the methods used for collecting the hemodynamic data that were analyzed in this study. A detailed description of all *in vivo* experimental procedures and animal preparations blood pressure and renal blood flow data acquisition from uninjured and SCI mice are described in our prior publication^23^. All *in vivo* experiments utilized adult female C57BL/6 mice, aged 2 to 6 months, obtained from Jackson Laboratories. Only female mice were used due to high-level SCI having more profound effects on renal function and structure^23^. Prior to any surgical interventions, the mice were housed in the institutional animal facility and allowed to acclimate for a minimum of one week. They were maintained in a temperature-controlled environment at 22°C under a standard 12:12-hour light-dark cycle, with *ad libitum* access to standard chow and water. All experimental protocols and animal care procedures strictly complied with the United States Animal Welfare Act and were formally approved by the Institutional Animal Care and Use Committee of Drexel University.

All hemodynamic data acquisition was performed using LabChart 8 software (ADInstruments, Colorado Springs, CO) and sampled at a rate of 1 kHz. All offline data analysis was conducted using custom-written software implemented in MATLAB (R2025a, The MathWorks, Inc., Natick, MA).

### Time Domain Analysis

Quantitative analysis was conducted on the blood pressure data in the time domain. For each signal, peak envelope detection was first performed to extract the systolic blood pressure (SBP) waveform, and a Savitzky-Golay filter with window size 50 was subsequently applied to smooth the signal. After determining the time index of the injection, the signal was segmented to remove the initial baseline period and commence at the time of the injection. Outliers were identified with the generalized extreme studentized deviate test and filled with the nearest non-outlier value. After pre-processing, an exponential decay curve of the form

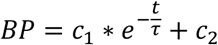

was fit to the signal, where *c_1_* and *c_2_* are the initial value relative to the asymptote and the asymptote, respectively, and τ is the time constant in seconds. A one-way ANOVA test was performed on the time constant across treatment groups. Tukey’s Honestly Significant Difference (HSD) test was used for post-hoc multiple comparisons between treatment groups with the control group, 24hr and 4wk samples within an injury group, and T3Tx and T10Tx samples at the same time post-injury. All comparisons with a *P*-value of < 0.05 were considered statistically significant.

### Frequency Domain Analysis

Due to significant variability in the macroscopic RBF waveforms across and within groups, particularly for samples with high-level injury, we were unable to determine a single model in the time domain that could be used as a viable measure of comparison. As such, RBF data was analyzed in the frequency domain, with analysis focused on the frequency range from 0.08 to 0.50 Hz, which encompasses the myogenic response and sympathetic vasomotor activity in mice^14,18,20,24^. For each subject, RBF data collected at resting state was used as a baseline for comparison. The baseline data was preprocessed with a Savitzky-Golay filter with window size 10 for smoothing. According to Welch’s method, the signal was divided into 32,768 sample segments with 50% overlap between adjacent segments to reduce noise and variance^13,18,25^. A Hanning window was applied to each segment to reduce spectral leakage^15,25,26^. The power spectral density (PSD) was calculated for each segment, scaled by the number of samples per segment, and averaged to yield an overall periodogram. The frequency range of interest was divided into 7 equal bins of size 0.06 Hz, and the area under the PSD curve in each of the 7 frequency bins was calculated and scaled by a correction factor equal to the average power of the Hanning window to determine the average signal power in the respective bins in accordance with Parseval’s Theorem.

RBF data measuring response to the injection was processed similarly, with the exception that the data segments were not averaged. PSD and area under the curve calculations for average signal power (with the same scaling factors noted previously) in the 7 frequency bins were performed for all segments separately to examine variation in the frequency domain over the duration of the signal. In addition, the power values were normalized relative to the baseline power in the corresponding frequency bin to quantify relative change and to facilitate comparison across samples. Relative power was calculated with the decibel signal power ratio by the formula

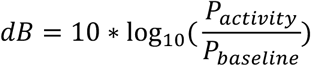

and was used for normalization to account for the power law of frequency band power^27^. For each subject, 4 segments were selected for further analysis – injection period (centered around the administration of the NE bolus), acute response (beginning directly after the injection), recovery period (beginning immediately following the end of the acute response period), and return to baseline (ending one-segment’s length from the end of the signal).

Two-way ANOVA tests were used for statistical analysis across treatment groups and frequency bins. One-way ANOVA tests were performed within each frequency bin to compare power values across treatment groups. Post-hoc analysis was conducted for all one-way ANOVA tests with a *P*-value < 0.10 using Tukey’s HSD test to evaluate differences between treatment groups and control, 24hr and 4wk samples within an injury group, and T3Tx and T10Tx samples at the same time post-injury. All comparisons with a *P*-value of < 0.05 were considered statistically significant.

## RESULTS

Representative waveforms for the raw BP and RBF data are shown in Figure 1 for response to NE administration in each of the 5 treatment groups (Control, T10Tx 24hr, T10Tx 4wk, T3Tx 24hr, and T3Tx 4wk). In uninjured mice, the increase in BP caused by a 10 µg/kg bolus intravenous administration of NE is associated with a sharp decrease in RBF, which is also observed in the T10Tx SCI mice both 24 hours and 4 weeks after injury. In contrast, the RBF in the T3Tx SCI mice increased alongside the increase in BP both 24 hours and 4 weeks after injury.

**Figure 1.**
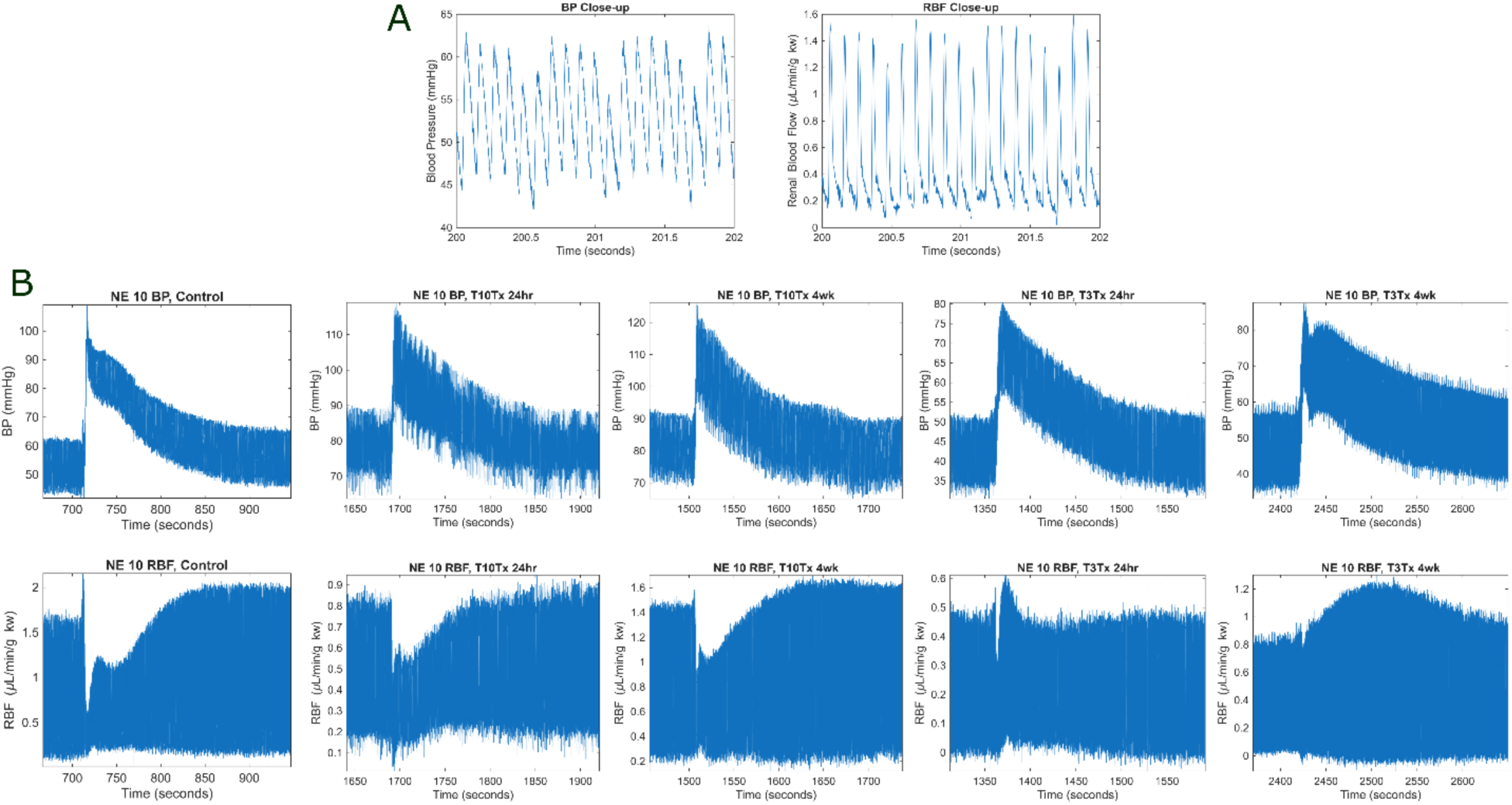
High thoracic-level spinal cord injury (SCI) disrupts renal vascular response to increases in systemic pressure. *A*: Close-up view of BP and RBF waveforms, which show periodic patterns around 10 Hz, corresponding to the cardiac cycle, and 2 Hz, identified to be a breathing artifact. *B*: Representative waveforms for blood pressure (BP) and renal blood flow (RBF) recovery response after a 10 µg/kg NE (norepinephrine) administration.

### Time Domain Analysis

Representative SBP waveforms and curve fittings are shown in Figure 2A for all treatment groups. The time constant of the exponential decay model, representing the rate of SBP recovery after a 10 µg/kg administration of norepinephrine, was compared across treatment groups. Out of 31 curves, 3 were excluded due to inadequacy of fit (R^2^ < 0.90) or abnormal measurements.

**Figure 2.**
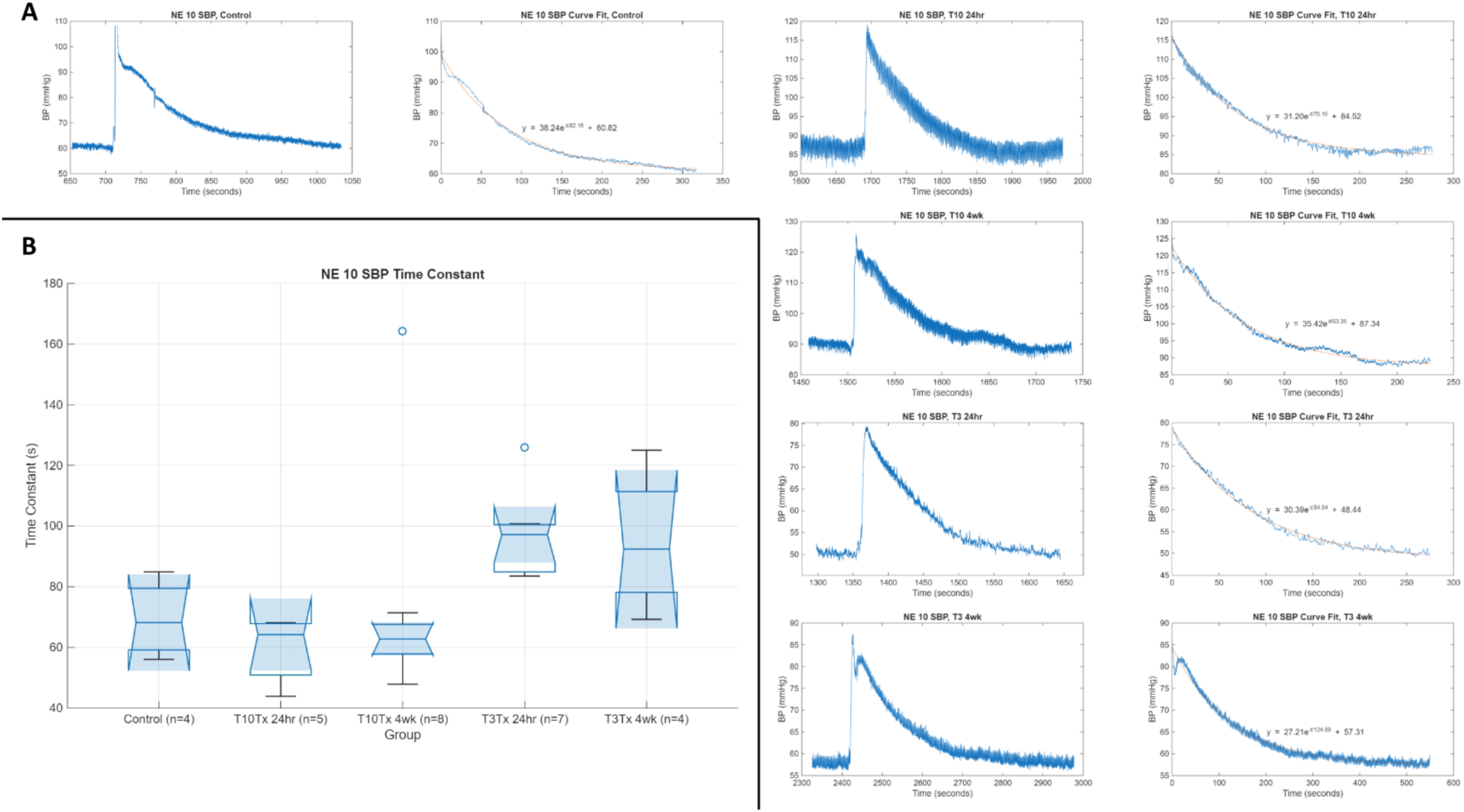
Slower rate of systolic blood pressure (SBP) recovery is associated with mice with spinal cord transection at thoracic level 3 (T3Tx). *A*: Representative SBP waveforms and curve fittings of the recovery response after a 10 µg/kg NE (norepinephrine) administration. *B*: Box chart of time constants (mean ± SE) in seconds for exponential decay curves fit to SBP signals across treatment groups.

A one-way ANOVA was performed to compare the effect of SCI on rate of SBP recovery after a 10 µg/kg administration of NE. The test revealed that there was no statistically significant difference in mean SBP recovery rate between at least two groups (F(4, 27) = 2.5887, p = 0.0636). Tukey’s HSD test found that there was no statistically significant difference in mean SBP recovery rate for any of the comparisons investigated. The results are shown in Figure 2B and Table 1.

**Table 1.**
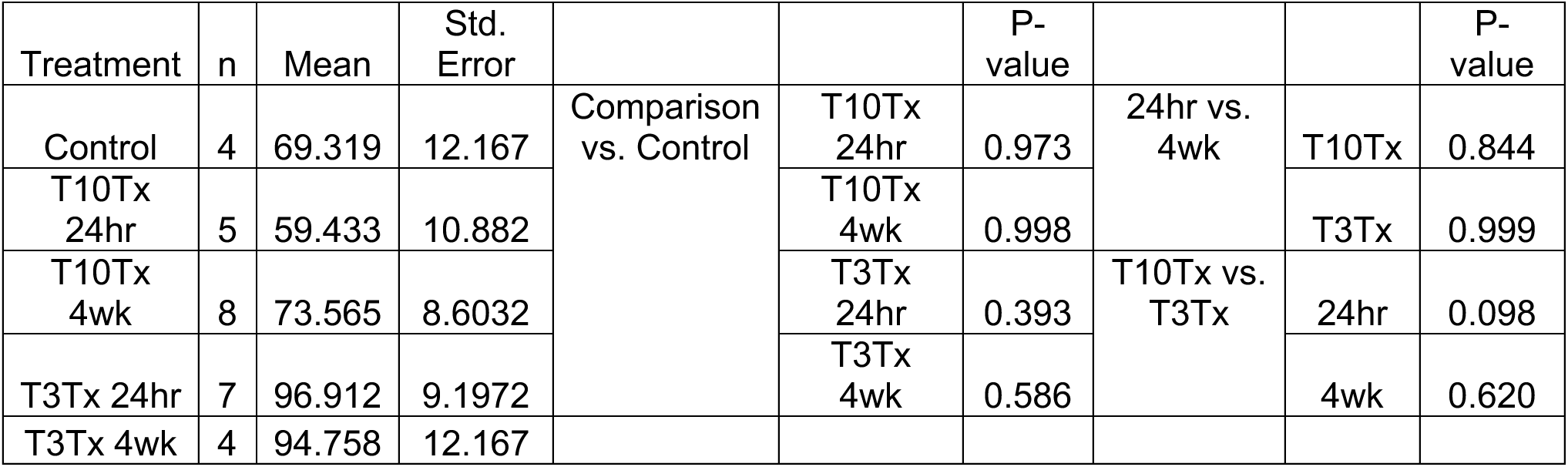
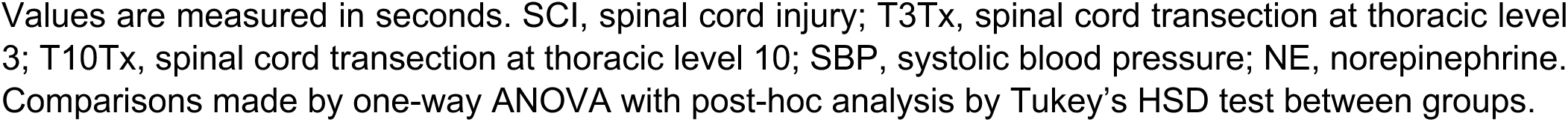
Effect of SCI in T10Tx and T3Tx female mice 24hr or 4wk after injury on rate of SBP recovery response after NE (10 µg/kg) administration.

The T3Tx 24hr group exhibited the largest mean time constant of 96.912, suggesting the slowest SBP recovery after the injection, while the T10 24hr group exhibited the smallest mean time constant of 59.433, suggesting the fastest SBP recovery after the injection. These differences are apparent in the observation that the T3Tx group had a P-value of 0.098 compared to the T10Tx group at 24 hours post-SCI. After 4 weeks, both groups exhibit stabilization towards the control group, particularly in the T10Tx group.

### Frequency Domain Analysis

Analysis on RBF signals was conducted over a range of 0.08 to 0.50 Hz. Representative normalized spectrograms showing the signal power in response to NE (10 µg/kg) administration relative to the subject baseline are shown in Figure 3 for all treatment groups. A representative RBF waveform in the control group with the 4 injection-recovery stages identified is shown in Figure 4B. Of the 31 samples, 1 sample was excluded due to measurement error.

**Figure 3.**
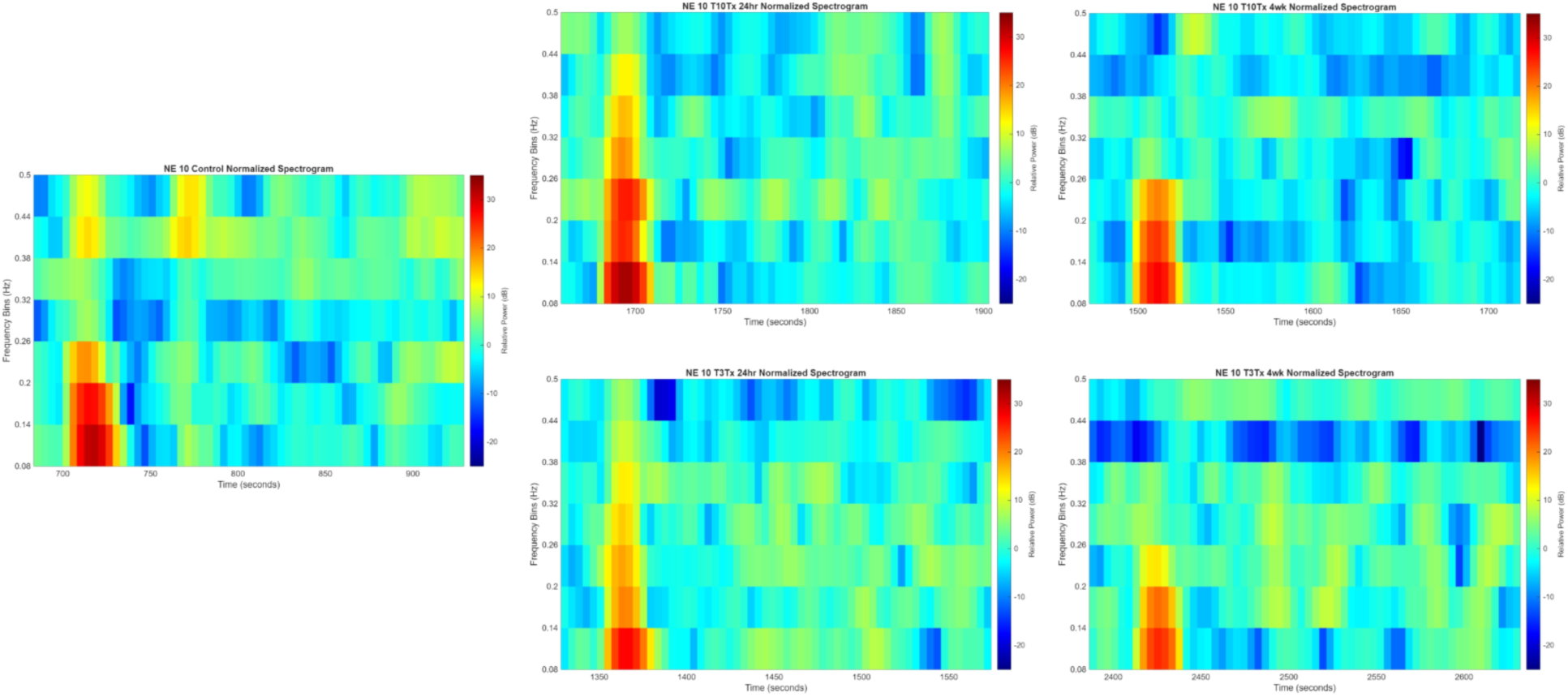
High-thoracic level spinal cord injury (SCI) causes reduced renal blood flow (RBF) activity in the frequency ranges associated with the myogenic response and sympathetic vasomotor activity. Representative spectrograms of the response to a 10 µg/kg NE (norepinephrine) administration in the 0.08 to 0.50 Hz range indicate that normalized signal power in the sympathetic range is severely attenuated in mice with spinal cord transection at thoracic level 3 (T3Tx), particularly 4 weeks after injury. Activity in the myogenic range is also reduced, both in magnitude and duration.

**Figure 4.**
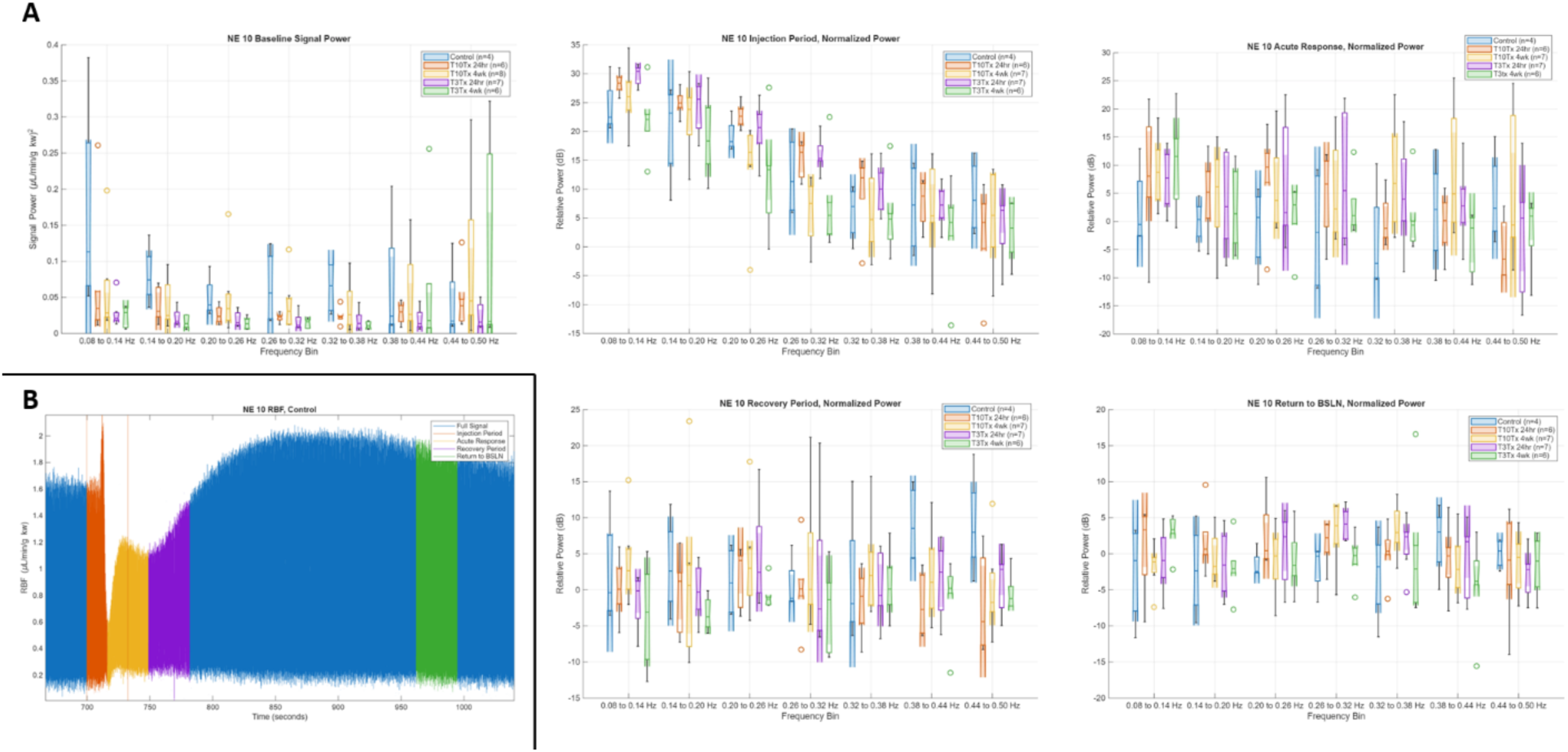
Reduced renal blood flow (RBF) activity in the myogenic and sympathetic ranges is observed in mice with spinal cord injury (SCI), with larger effects observed in those with spinal cord transection at thoracic level 3 (T3Tx). *A*: Box charts of signal power (mean ± SE) in (µL/min/g kw)^2^ for baseline activity or relative power (mean ± SE) in dB for 4 injection-recovery stages across treatment groups and frequency bins of size 0.06 Hz from 0.08 to 0.50 Hz. *B*: Representative control RBF waveform with the 4 injection-recovery stages (injection, acute response, recovery, and return to baseline) identified.

A two-way ANOVA was performed to compare the effect of SCI injury on rate of SBP recovery after a 10 µg/kg administration of NE for the baseline power and each of the 4 stages. One-way ANOVA tests were performed within each frequency bin to compare power values across treatment groups, with post-hoc analysis using Tukey’s HSD test conducted for all one-way tests with a P-value < 0.10. The results are shown in Figure 4A.

At baseline, a two-way ANOVA test revealed that there was not a statistically significant interaction between the effects of SCI injury and frequency range (F(24, 182) = 1.072, p = 0.380). Simple main effects analysis showed that SCI injury had a statistically significant effect on mean signal power (F(4, 182) = 5.632, p < 0.001) and that frequency range had a statistically significant effect on mean signal power (F(6, 182) = 2.542, p = 0.022). One-way ANOVA tests revealed that there was a statistically significant difference in mean signal power between at least two groups in the 0.14 to 0.20 Hz, 0.26 to 0.32 Hz, and 0.32 to 0.38 Hz frequency ranges. Tukey’s HSD indicated that these differences were in the T3Tx group at both 24 hr and 4 wk after injury with respect to the control group, as seen in Table 2.

**Table 2.**
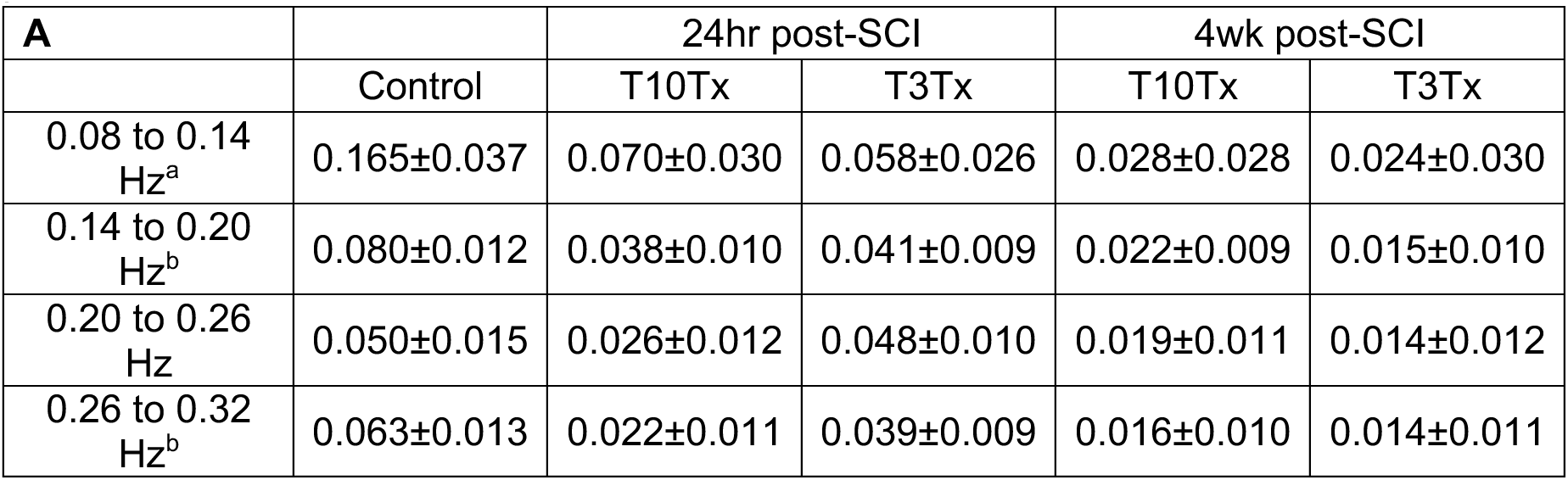
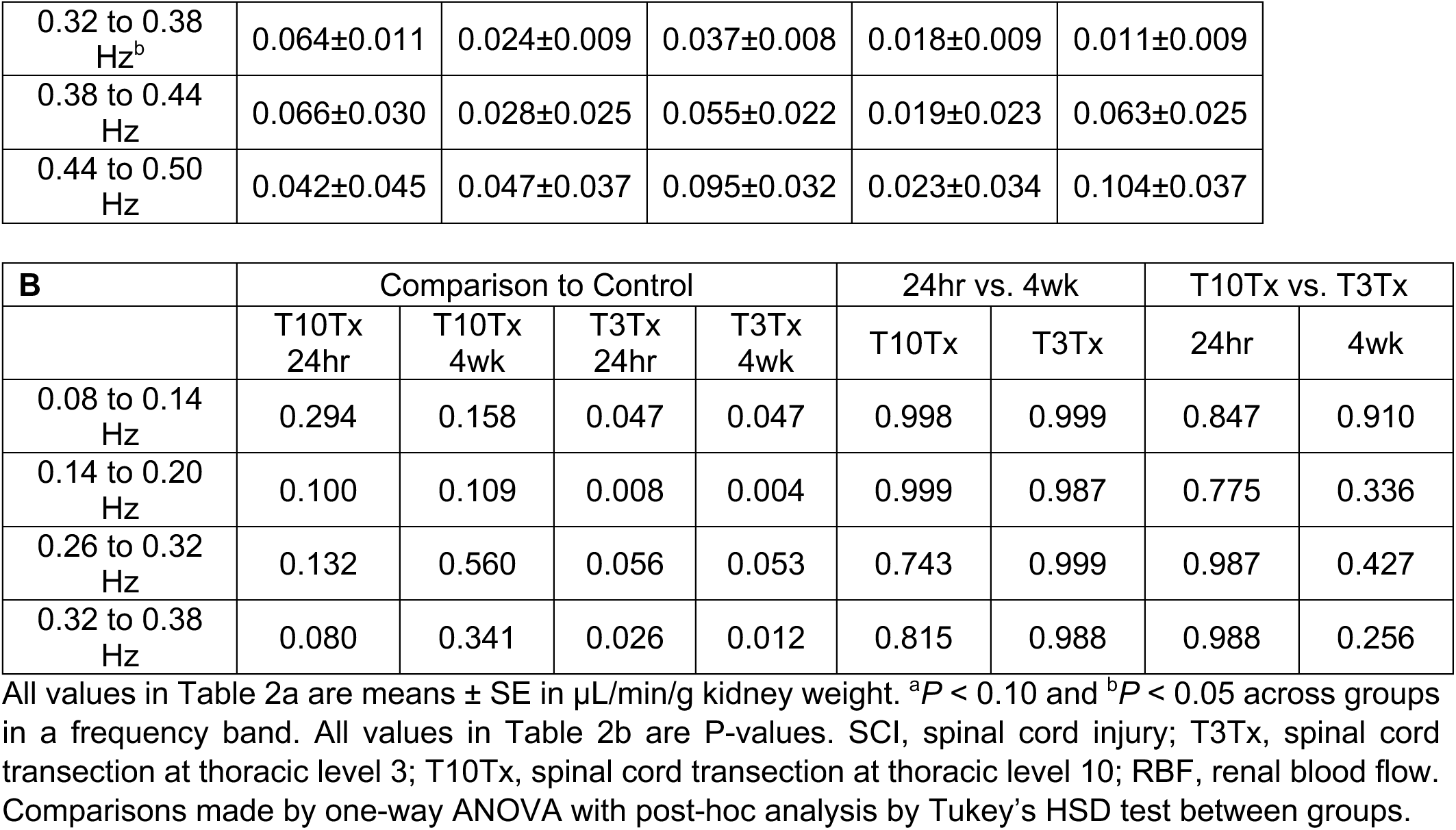
Effect of SCI in T10Tx and T3Tx female mice 24hr or 4wk after injury on baseline RBF signal power.

The injection period was defined as the time segment during which the 10 µg/kg bolus of norepinephrine was administered. A two-way ANOVA test revealed that there was not a statistically significant interaction between the effects of SCI injury and frequency range (F(24, 175) = 0.828, p = 0.698). However, simple main effects analysis showed that SCI injury had a statistically significant effect on mean relative power (F(4, 175) = 7.358, p < 0.001) and that frequency range had a statistically significant effect on mean relative power (F(6, 175) = 57.483, p < 0.001). One-way ANOVA tests revealed that there was a statistically significant difference in mean relative power between at least two groups in the 0.08 to 0.14 Hz and 0.26 to 0.32 Hz frequency ranges, with the largest difference identified by Tukey’s HSD test to be between 24 hr and 4 wk post-SCI for the T3Tx group. Results are summarized in Table 3.

**Table 3.**
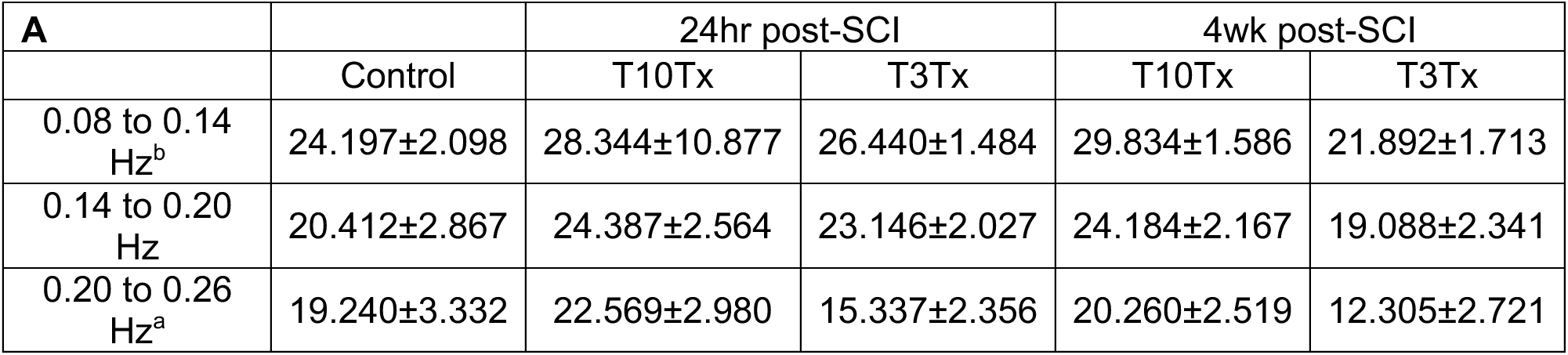
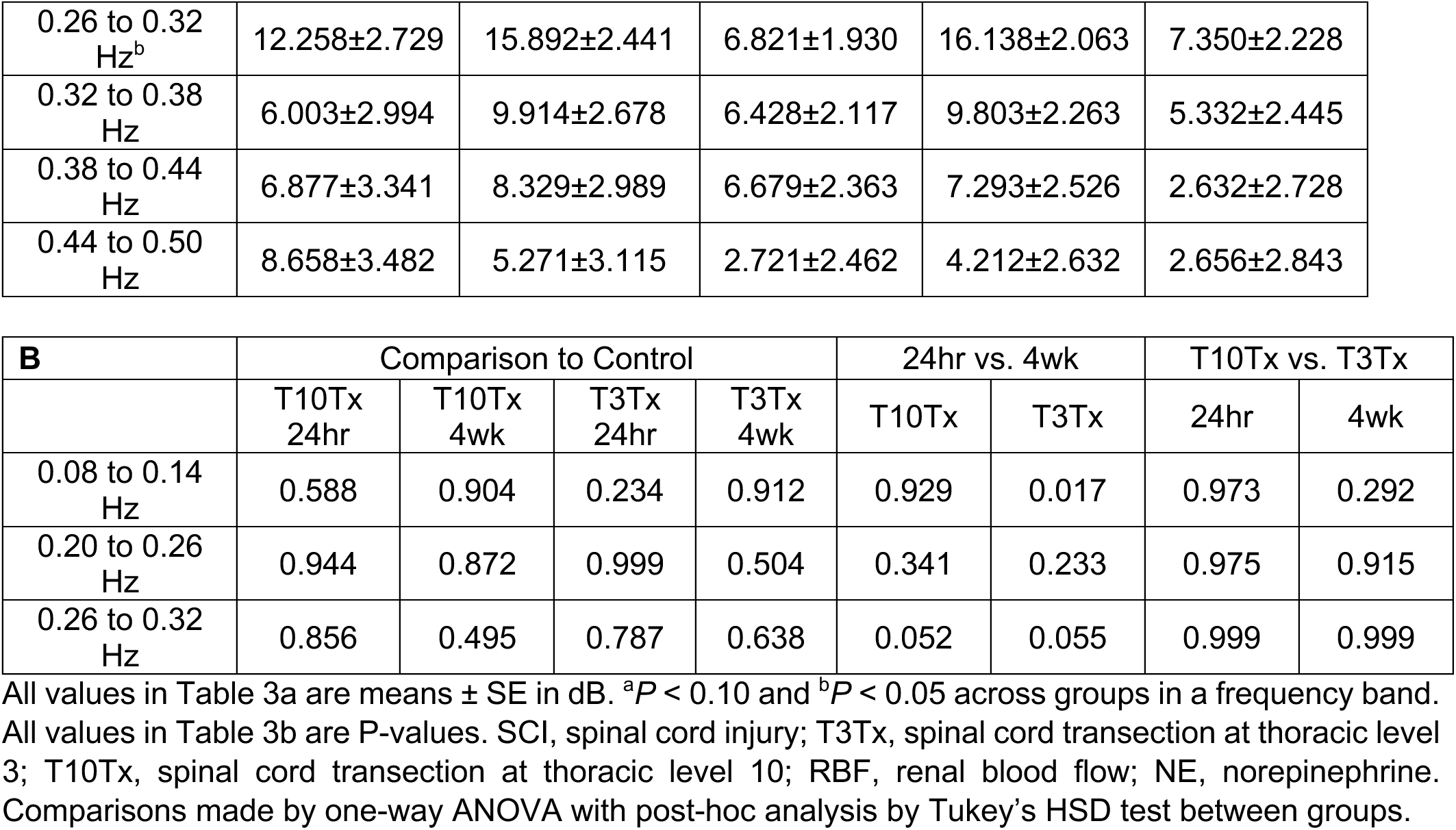
Effect of SCI in T10Tx and T3Tx female mice 24hr or 4wk after injury on relative RBF signal power during period of NE (10 µg/kg) administration.

The acute response period was defined as the time segment beginning directly following the 10 µg/kg administration of norepinephrine. A two-way ANOVA test revealed no statistically significant interaction between the effects of SCI injury and frequency range (F(24, 175) = 0.812, p = 0.719). However, simple main effects analysis showed that SCI injury had a statistically significant effect on mean relative power (F(4, 175) = 3.681, p = 0.007) and that frequency range had a statistically significant effect on mean relative power (F(6, 175) = 2.264, p = 0.040). One-way ANOVA tests revealed that there was no statistically significant difference in mean relative power between at least two groups in any of the frequency ranges, as seen in Table 4. As such, post-hoc tests were not performed.

**Table 4.** Effect of SCI in T10Tx and T3Tx female mice 24hr or 4wk after injury on relative RBF signal power during period of acute response after NE (10 µg/kg) administration.

|  | Control | 24hr post-SCI |  | 4wk post-SCI |  |
| --- | --- | --- | --- | --- | --- |
|  |  | T10Tx | T3Tx | T10Tx | T3Tx |
| 0.08 to 0.14 Hz | 2.326±3.801 | 11.800±3.400 | 6.419±2.688 | 7.602±2.873 | 10.574±3.104 |
| 0.14 to 0.20 Hz | -<br>0.017±3.714 | 6.663±3.322 | 2.833±2.626 | 3.981±2.808 | 2.214±3.033 |
| 0.20 to 0.26 Hz | -<br>1.001±4.017 | 11.109±3.593 | 4.236±2.841 | 7.303±3.037 | 1.016±3.280 |
| 0.26 to 0.32 Hz | -<br>1.634±4.432 | 6.993±3.964 | 3.718±3.134 | 7.791±3.350 | 2.732±3.619 |
| 0.32 to 0.38 Hz | -<br>3.743±4.028 | 0.820±3.603 | 6.755±2.848 | 4.863±3.045 | 0.557±3.289 |
| 0.38 to 0.44<br>Hz | 1.642±2.098 | -0.219±1.877 | 7.402±1.484 | 3.015±1.586 | -3.441±1.713 |
| 0.44 to 0.50<br>Hz | 4.055±4.914 | -4.094±4.696 | 3.917±3.475 | -<br>2.522±3.715 | -1.669±4.013 |

All values are means ± SE in dB. SCI, spinal cord injury; T3Tx, spinal cord transection at thoracic level 3; T10Tx, spinal cord transection at thoracic level 10; RBF, renal blood flow; NE, norepinephrine. Comparisons made by one-way ANOVA test.

The recovery period was defined as the time segment beginning immediately following the end of the time segment previously identified as the acute response period. The two-way ANOVA test revealed that there was not a statistically significant interaction between the effects of SCI injury and frequency range (F(24, 175) = 0.732, p = 0.814). Simple main effects analysis showed that SCI injury had a statistically significant effect on mean relative power (F(4, 175) = 4.830, p = 0.001), but frequency range did not have a statistically significant effect on mean relative power (F(6, 175) = 0.520, p = 0.814). One-way ANOVA tests revealed that there was a statistically significant difference in mean relative power between at least two groups in the 0.38 to 0.44 Hz frequency range. Tukey’s HSD test indicated that notable differences were in the T10Tx 24hr and T3Tx 4wk groups, as see in Table 5.

**Table 5.**
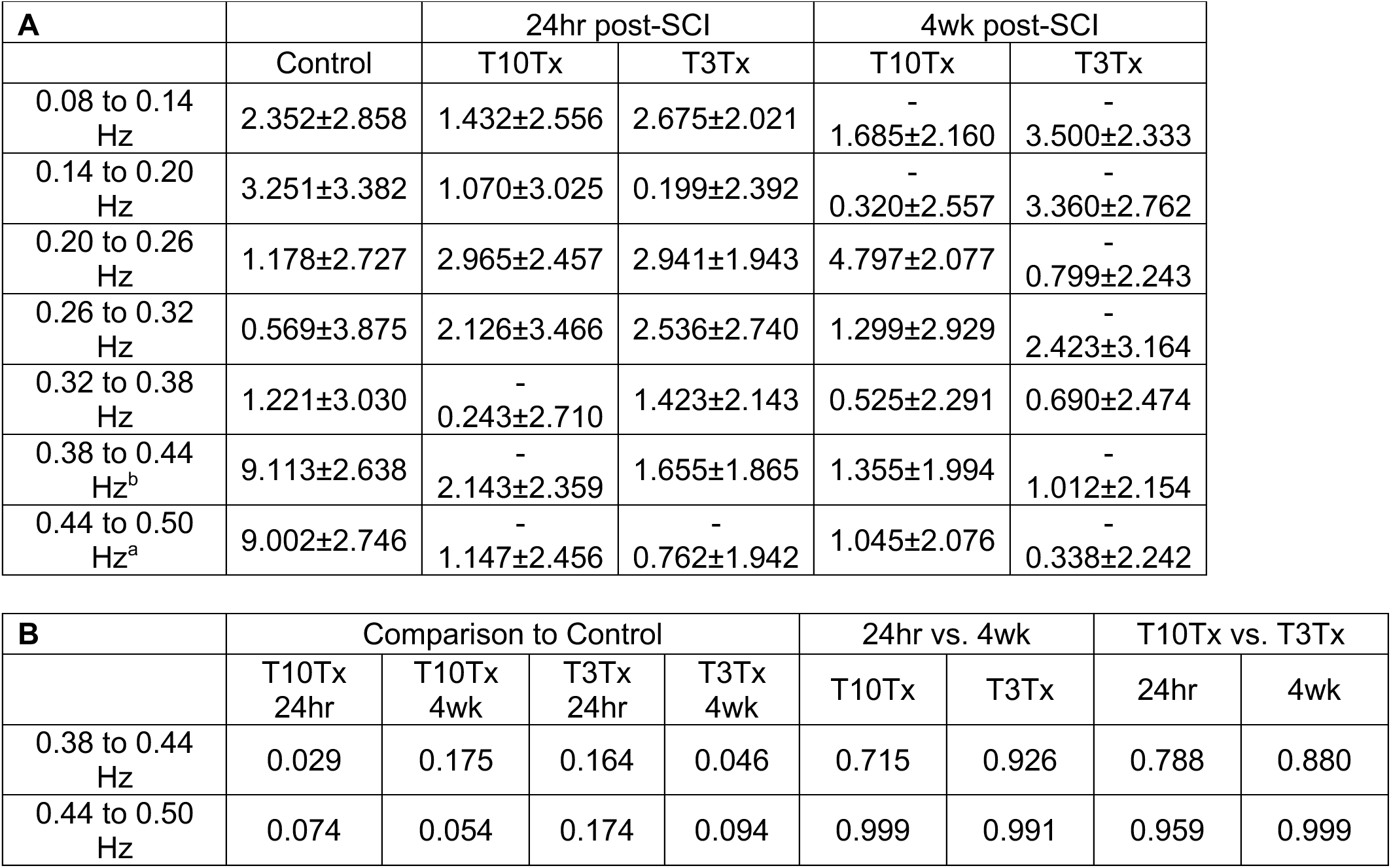
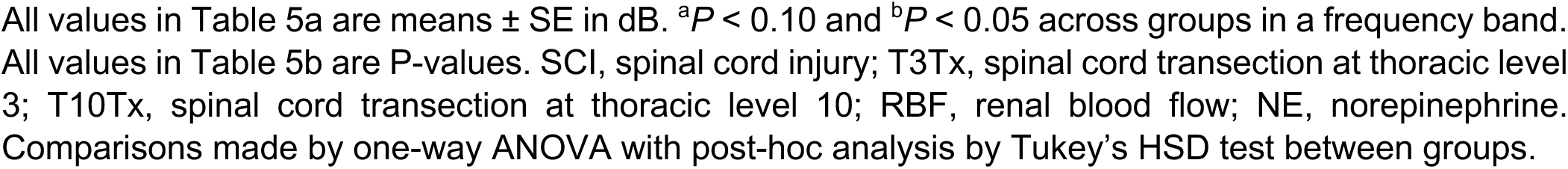
Effect of SCI in T10Tx and T3Tx female mice 24hr or 4wk after injury on relative RBF signal power during recovery period following NE (10 µg/kg) administration.

The RBF was determined to have returned to the baseline when the upper envelope of the RBF signal returned to pre-injection levels. A representative segment ending at minimum one-segment’s length from the end of the signal without outlying spikes was chosen for this analysis. A two-way ANOVA test revealed that there was not a statistically significant interaction between the effects of SCI injury and frequency range (F(24, 175) = 1.015, p = 0.449). Simple main effects analysis showed that SCI injury did not have a statistically significant effect on mean relative power (F(4, 175) = 1.065, p = 0.376). Frequency range also did not have a statistically significant effect on mean relative power (F(6, 175) = 1.273, p = 0.272). One-way ANOVA tests revealed that there was a statistically significant difference in mean relative power between at least two groups in the 0.26 to 0.32 Hz range, but post-hoc Tukey’s HSD analysis yielded no statistically significant results, as seen in Table 6.

**Table 6.**
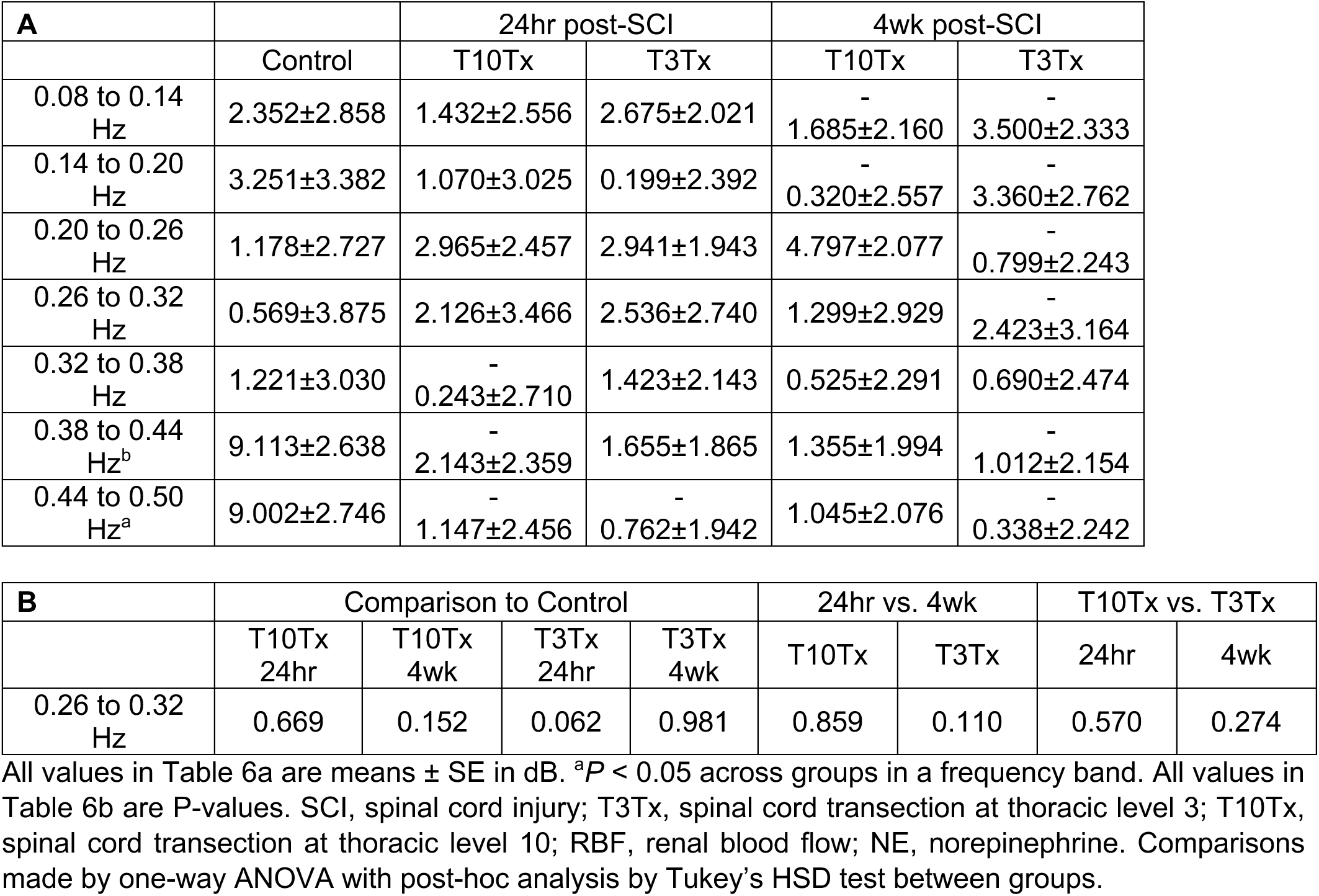
Effect of SCI in T10Tx and T3Tx female mice 24hr or 4wk after injury on relative RBF signal power after return to baseline.

## DISCUSSION

The present study demonstrates that spinal cord injury (SCI) level and chronicity exert profound effects on renal hemodynamic regulation by disrupting supraspinal autonomic control of the renal vasculature. Using time- and frequency-domain analyses of in vivo blood pressure and renal blood flow, we show that high-thoracic SCI produces a persistent impairment of dynamic renal autoregulation, characterized by paradoxical renal hyperperfusion during acute hypertensive stimuli and sustained loss of myogenic and sympathetic vasomotor activity. These alterations emerge acutely following injury and are exacerbated during the chronic phase, consistent with progressive autonomic-vascular uncoupling after neurotrauma. In contrast, low-thoracic SCI preserves critical descending sympathetic pathways, allowing partial recovery of renal vasomotor control and stabilization of renal hemodynamics over time.

### Autonomic disruption as a driver of renal dysregulation after SCI

Disruption of supraspinal autonomic pathways is a hallmark consequence of SCI and a major contributor to cardiovascular instability following neurotrauma^1–3^. High-thoracic injuries interrupt descending sympathetic control to the splanchnic and renal vascular beds, producing an acute loss of vasomotor tone that manifests clinically as neurogenic shock, hypotension, and reduced organ perfusion^2,21^. Our findings extend this framework by demonstrating that high-thoracic SCI not only alters systemic blood pressure regulation but also fundamentally disrupts the kidney’s ability to dynamically buffer pressure changes, thereby exposing the renal microvasculature to injurious hemodynamic stress.

In the chronic phase of high-level SCI, maladaptive plasticity within decentralized spinal autonomic circuits promotes exaggerated sympathetic reflexes and extreme blood pressure variability, particularly during episodes of autonomic dysreflexia^3,5,6,22^. The paradoxical increases in renal blood flow observed during norepinephrine-induced hypertension in T3-injured mice are consistent with a loss of coordinated vasomotor control and suggest dissociation between systemic pressure regulation and local renal vascular responses. This phenomenon likely contributes to the progressive renal injury observed clinically and experimentally following high-level SCI^7,9,28^.

### Frequency-specific loss of renal vasomotor control

Dynamic renal autoregulation depends on tightly coordinated interactions between myogenic, tubuloglomerular, and neurally mediated mechanisms that operate across distinct frequency ranges^12–14,17–19^. Frequency-domain analysis of renal blood flow (RBF) provides a powerful approach to resolving these components and has been used to identify myogenic and sympathetic contributions to renal vasomotion under physiological and pathological conditions^17–20^. Applying this framework to SCI, we demonstrate that high-thoracic injury produces a broadband reduction in RBF signal power within frequency ranges associated with both myogenic (0.08–0.20 Hz) and sympathetic (0.20–0.38 Hz) activity.

These findings indicate that loss of supraspinal autonomic input disrupts not only neural vasomotor signaling but also the intrinsic myogenic responsiveness of the renal microvasculature. This impairment likely reflects a combination of altered vascular smooth muscle function and structural remodeling of resistance vessels, as previously described following chronic SCI^23^. In contrast, low-thoracic SCI largely preserved baseline spectral activity and demonstrated recovery of frequency-specific vasomotor control over time, consistent with sparing of descending sympathetic projections to the renal bed^1,21^.

### Temporal evolution of renal autoregulatory failure

The differential effects observed between acute and chronic phases of SCI highlight the evolving nature of autonomic and vascular dysfunction after neurotrauma. In the acute phase, exaggerated vascular responses to sympathetic stimulation in high-thoracic SCI may reflect adrenergic receptor hypersensitivity following sudden withdrawal of tonic sympathetic drive^21^. Over time, however, chronic exposure to extreme blood pressure variability and maladaptive autonomic reflexes likely drives progressive vascular remodeling, resulting in a sustained loss of dynamic autoregulatory capacity^3,23^.

Our time-domain analyses of systolic blood pressure recovery support this interpretation, revealing delayed pressure normalization following sympathetic stimulation in high-thoracic SCI, particularly during the acute phase. Although some stabilization was observed chronically, this apparent recovery of systemic pressure control was not accompanied by restoration of normal renal vasomotor dynamics, underscoring the dissociation between systemic and organ-specific hemodynamic regulation after SCI.

### Implications for secondary organ injury after neurotrauma

Secondary injury processes are increasingly recognized as major determinants of long-term outcomes after SCI^3,4^. While much attention has focused on neuroinflammation, vascular dysfunction, and metabolic changes within the injured spinal cord, comparatively less emphasis has been placed on injury to peripheral organs that arises from autonomic dysregulation. The kidney is uniquely susceptible to such injury due to its high perfusion demand and reliance on stable arterial pressure to protect the microcirculation^7,10,12^.

Our findings provide mechanistic insight into how SCI-induced autonomic disruption compromises renal autoregulation, offering a potential explanation for the increased incidence of renal dysfunction and chronic kidney disease observed in individuals with high-level SCI^7,9,28^. By identifying frequency-specific loss of vasomotor control, this work suggests that renal injury after SCI is not solely a consequence of hypotension or hypertension per se, but rather of impaired dynamic buffering of hemodynamic stress.

### Limitations and future directions

Several limitations should be considered when interpreting these findings. First, sympathetic activity was inferred from frequency-domain characteristics of RBF rather than measured directly. While this approach is well supported by prior experimental and theoretical work^17–20^, future studies combining direct sympathetic nerve recordings with frequency-domain analysis would further strengthen mechanistic interpretation. Second, this study focused on female mice, reflecting evidence that high-level SCI produces more pronounced renal dysfunction in females^23^; whether similar dynamics occur in males remains to be determined. Finally, structural correlates of the observed functional changes were not directly assessed in this study but are strongly supported by prior histological analyses following chronic SCI^23^.

## Conclusions

In summary, this study demonstrates that spinal cord injury (SCI) level and chronicity critically determine renal hemodynamic regulation through disruption of supraspinal autonomic control. Using time- and frequency-domain analyses, we show that high-thoracic SCI produces a persistent, broadband impairment of dynamic renal autoregulation, characterized by loss of myogenic and sympathetic vasomotor activity and paradoxical renal blood flow responses to acute hypertensive stimuli. These deficits emerge early after injury and persist chronically, consistent with progressive autonomic-vascular uncoupling following neurotrauma. In contrast, low-thoracic SCI preserves key descending sympathetic pathways, maintaining baseline renal vasomotor activity and permitting recovery of dynamic autoregulatory responses over time.

From a clinical perspective, these findings suggest that renal vulnerability after high-level SCI may persist despite apparent stabilization of systemic blood pressure. Blood pressure variability and episodic autonomic disturbances characteristic of high-thoracic SCI may therefore expose the renal microvasculature to injurious hemodynamic stress that is not captured by conventional cardiovascular metrics. By linking autonomic disruption to frequency-specific impairment of renal autoregulation, this work provides a mechanistic framework for understanding secondary renal injury after SCI and highlights dynamic vascular control as a potential target for mitigating long-term organ dysfunction following neurotrauma.

## DATA AVAILABILITY

All source data (raw blood pressure and renal blood flow) used in this study are readily available upon reasonable request to the corresponding author.

## ACKNOWLEDGEMENTS

We thank members of the Osei-Owusu and Tom laboratories for technical support. The Visual Abstract was generated using Google NotebookLM.

## GRANTS

This work was supported by Craig H. Neilsen Foundation Grant 382566 to V.J.T. and P.O.; National Institutes of Health Grants R01NS1069080, R01 NS111761, R01NS085426, and R01NS122371 to V.J.T.; R01HL139754, R56DK132859-0A1 and R01HL174004-01 to P.O.

## DISCLOSURES

The authors have no conflict of interest, financial or otherwise, to disclose.

## AUTHOR CONTRIBUTIONS

PO, VJT, and EGC conceived and designed the research; VJT and PO performed experiments; AT, GK, EGC, and PO analyzed data, interpreted results of experiments, prepared figures, drafted manuscript, edited and revised manuscript; all authors approved the final version of the manuscript.

## Notes

### Competing Interest Statement

The authors have declared no competing interest.

